# Lack of evidence for the contribution of cone photoreceptors to human melatonin suppression and alerting response to light at night

**DOI:** 10.1101/2024.10.10.617409

**Authors:** Fatemeh Fazlali, Rafael Lazar, Faady Yahya, Christian Epple, Manuel Spitschan, Oliver Stefani, Christian Cajochen

## Abstract

Light exposure at night can suppress melatonin production and increase alertness, primarily through the action of melanopsin-containing intrinsically photosensitive retinal ganglion cells (ipRGCs). This study investigated whether cone photoreceptors also influence melatonin suppression and subjective alertness using non-visual metameric light emitted from a display. Forty-eight participants with normal trichromatic vision were exposed to three lighting conditions: a baseline (9 lx_mEDI_), constant background (149 lx_mEDI_), and cone-modulated flickering light targeting different cone combinations and post-receptoral channels (149 lx_mEDI_) for 2 hours after their habitual bedtime. Salivary melatonin levels and subjective alertness were measured throughout a 9-h protocol. Bayesian analysis showed that cone-modulated flickering light did not significantly affect melatonin suppression or alertness, providing evidence against the hypothesis that cone photoreceptors contribute to these non-visual effects of light. In conclusion, our results suggest cone photoreceptors do not play a measurable role in light’s effects on melatonin suppression and subjective alertness at night.

## Introduction

Ambient light of varying wavelengths from different directions enters the human eye, passes through the pupil and lens, and reaches the retina. The retina is a critical sensory organ that mediates both the image and non-image forming effects of light^1–3^. In humans, the retina comprises classical photoreceptors, rods^4^ and cones (S-, M-, L-cone)^5,6^, which have different spectral sensitivities and non-classical melanopsin-containing intrinsically photosensitive retinal ganglion cells (ipRGCs)^7–9^ with peak sensitivity at around 480 nm^10^. These photoreceptors convert light into neural signals that are transmitted via the optic nerve to various brain regions^3,11^. Image formation is facilitated by signal transmission to the visual cortex via the thalamic lateral geniculate nucleus (LGN)^12–15^, while non-visual responses such as circadian regulation are mediated by the retinohypothalamic pathway, which projects primarily to the circadian pacemaker located in the suprachiasmatic nuclei (SCN)^16–18^.

Non-visual effects of light, including melatonin suppression are mediated predominantly by ipRGCs^19–22^. These cells play a crucial role in synchronising the internal circadian clock with the external light-dark cycle^23,24^. In visually intact humans, ipRGCs receive synaptic input from L-and M-cones and rods, and inhibitory input from S-cones^14,25^. While the role of ipRGCs in circadian regulation is well established, the specific contributions of retinal cones to these processes are less well understood. Evidence from animal studies suggests that cones can influence circadian responses^26–30^, but their exact role in human circadian physiology and their potential contribution to neuroendocrine regulation remains unclear. Previous research has shown that cones can influence pupillary light responses^31–34^, which are an indirect measure of circadian and neuroendocrine effects. However, it is not clear how cones contribute to the effects of light on human circadian physiology, particularly in terms of melatonin suppression and subjective alertness. Furthermore, it is not known which type of cone photoreceptor is most effective in mediating these circadian responses, or how these mechanisms operate under real-world lighting conditions.

In normal trichromatic humans, exposure to broadband light activates all types of retinal photoreceptors. To isolate the specific role of cones in circadian neuroendocrine responses, we used the silent substitution method^35^, which allows selective modulation of individual photoreceptor classes. By carefully tuning the spectral composition of the light emitted by LEDs, this method allows maximal stimulation of a target photoreceptor class while keeping the stimulation of other photoreceptors constant. This approach is particularly useful for assessing the contribution of post-receptoral channels^36–38^ in the suppression of nocturnal melatonin secretion by light.

In addition, each class of photoreceptors has different temporal response characteristics to time-varying light stimuli^39–42^, such as sinusoidal flicker or pulse trains. Flickering light stimuli can prevent cones from saturating and adapting^43–45^ to continuous light, potentially leading to stronger circadian and neuroendocrine effects compared to steady illumination. Studies have shown that flickering light, particularly at high frequencies, can induce more pronounced changes in retinal vessel diameter^46^, enhance visual cortex activation^42^, and reduce melatonin concentrations^41^ more effectively than continuous light under certain conditions. These findings suggest that flickering light could be a powerful tool for modulating circadian physiology and influencing alertness.

Here, we aimed to investigate the mechanistic role of cone photoreceptors in human neuroendocrine physiology during the evening. Specifically, we examined the effects of cone- modulated flickering light emitted from a non-visual metameric display on human melatonin suppression and subjective alertness. Using silent substitution with a custom-modulated five-primary display, we generated flickering stimuli that selectively activated single or multiple cone types with maximal contrast. Our primary outcome was the melatonin area under the curve (AUC), and the secondary outcome was subjective alertness in three lighting conditions, baseline control, constant background, and cone-modulated flickering light. We hypothesised that cone-modulated flickering light decreases melatonin production and increases subjective alertness compared to constant background light^41,42,47^. This study aims to provide new insights into the specific contributions of cone photoreceptors to neuroendocrine regulation in the human retina.

## Results

Forty-eight healthy participants with normal trichromatic vision (18-35 years, mean age = 24.8 ± 3.5 years, 50% female) were recruited and completed the study. In a 9-hour within-subject study protocol, participants were randomly exposed to three different experimental lighting conditions for 2 hours after their habitual bedtime (HBT) (**Figure 1**). A comprehensive overview of the irradiance, luminance, correlated colour temperature (CCT) and photoreceptor excitation levels in each condition can be found in **Table 1**. In all analyses, we report BF10, which represents the likelihood of the data under H1 compared to H0. Detailed interpretation of the BFs^48^, can be found in **Supplementary Table 1**.

**Figure 1:**
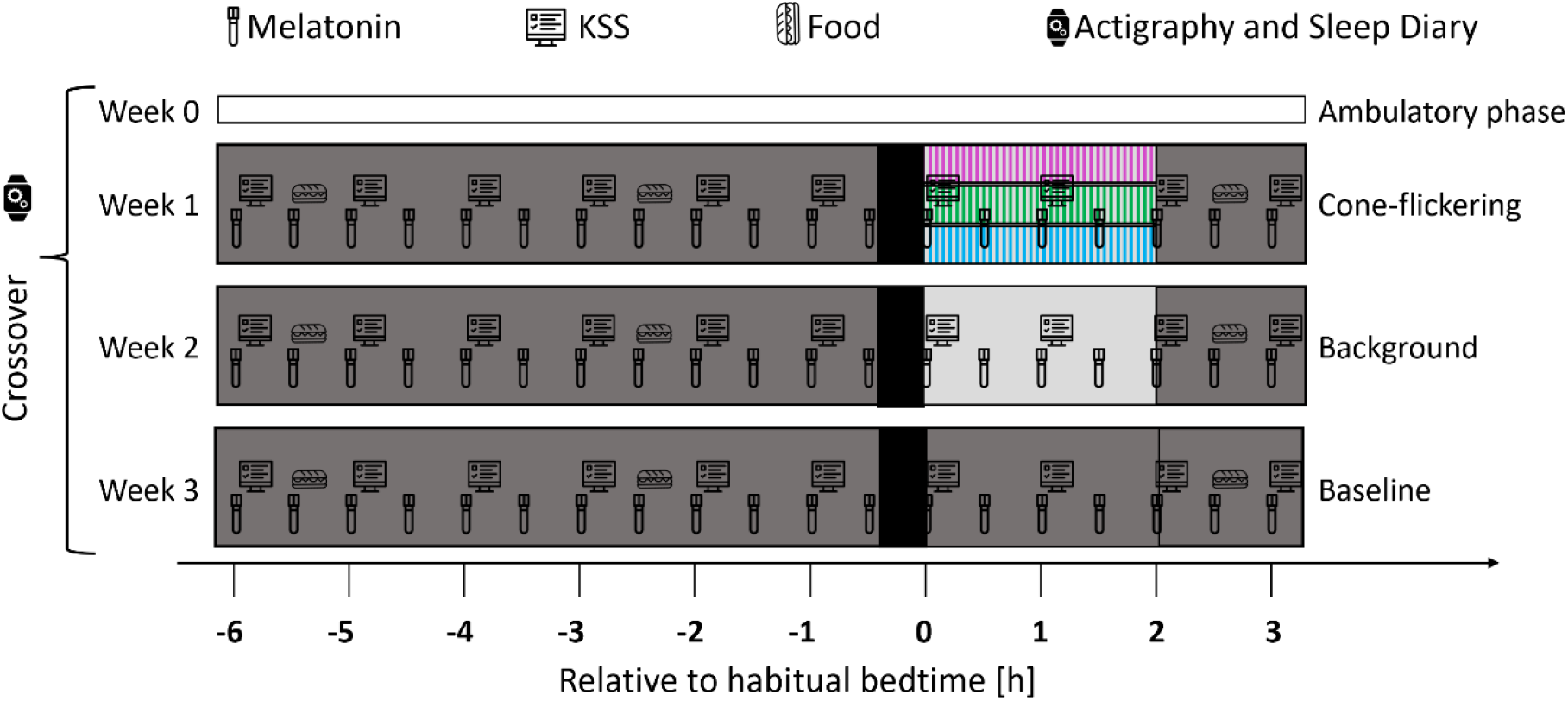
Whitin-subject study protocol. Participants underwent a 9-hour laboratory protocol starting 6 hours before and ending 3 hours after their habitual bedtime (HBT). The protocol consisted of three different light conditions: (1) a dim light baseline session (8 lx, 9 lx), (2) a constant background session (100 lx, 149 lx) for 2 hours, and (3) a cone-modulated flicker session (149 lx) for 2 hours. Participants were randomly assigned to one of the study sessions and also to one of the three cone-modulated light conditions: S-cone (blue), M-L (green) or S+M+L (magenta). They underwent a 20-minute dark-adaptation period prior to light exposure and a dim-light adaptation period prior to dark-adaptation and after light exposure. Salivary melatonin samples were collected every 30 minutes and subjective alertness scores (KSS) were recorded every hour. Participants were provided with 3 equicaloric sandwiches during the session to ensure consistent energy intake, which were given at specific intervals. During the ambulatory phase (>5 days) prior to the laboratory visits, participants maintained a regular sleep-wake schedule monitored by wrist actigraphy and an online sleep diary to ensure compliance with the study conditions.

**Table 1:**
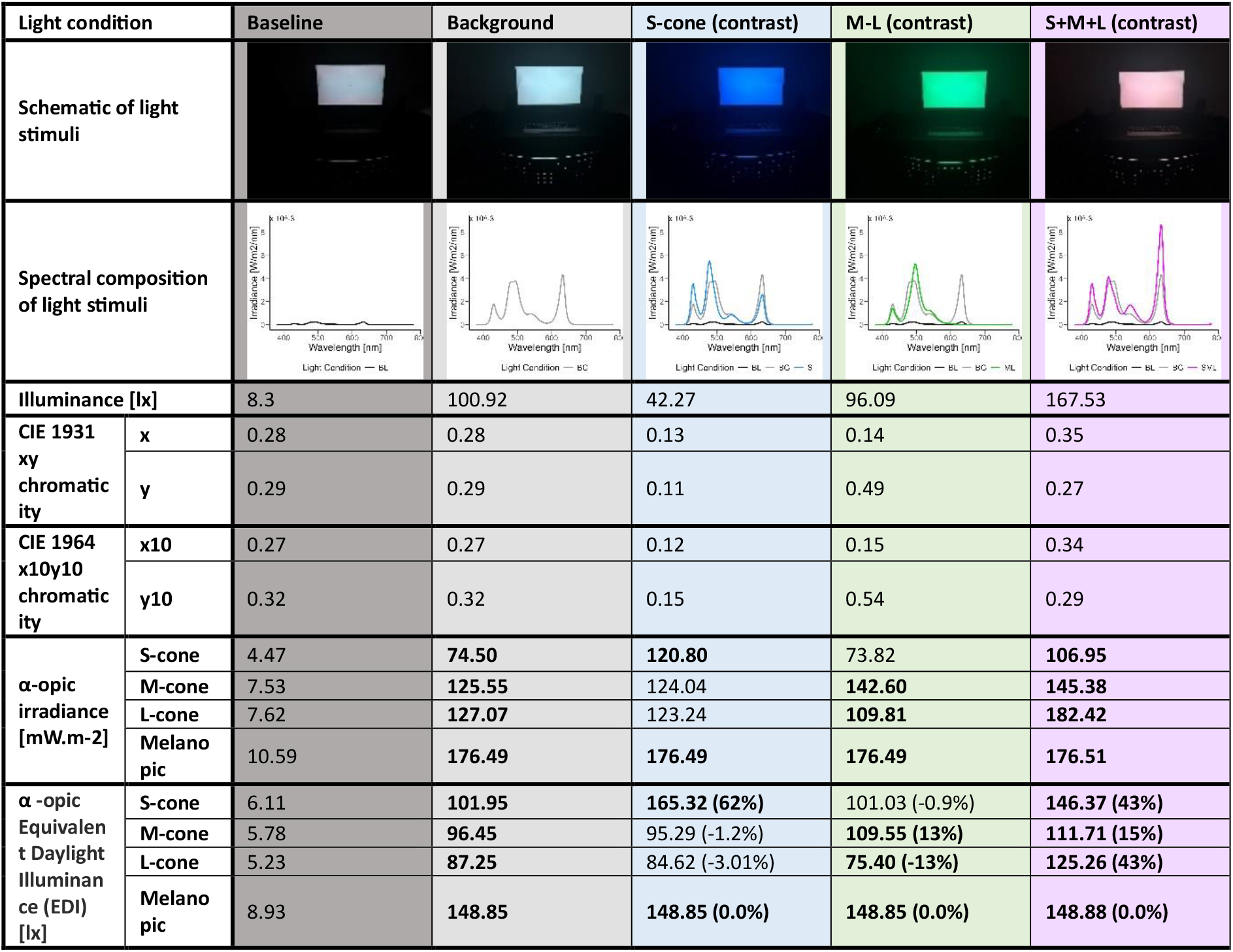
Overview light measurement of the experimental light conditions

| Light condition |  | Baseline | Background | S-cone (contrast) | M-L (contrast) | S+M+L (contrast) |
| --- | --- | --- | --- | --- | --- | --- |
| Schematic of light stimuli |  |  |  |  |  |  |
| Spectral composition of light stimuli |  |  |  |  |  |  |
| Illuminance [lx] |  | 8.3 | 100.92 | 42.27 | 96.09 | 167.53 |
| CIE 1931 xy chromaticity | x | 0.28 | 0.28 | 0.13 | 0.14 | 0.35 |
|  | y | 0.29 | 0.29 | 0.11 | 0.49 | 0.27 |
| CIE 1964 x10y10 chromaticity | x10 | 0.27 | 0.27 | 0.12 | 0.15 | 0.34 |
|  | y10 | 0.32 | 0.32 | 0.15 | 0.54 | 0.29 |
| α-opic irradiance [mW.m-2] | S-cone | 4.47 | 74.50 | 120.80 | 73.82 | 106.95 |
|  | M-cone | 7.53 | 125.55 | 124.04 | 142.60 | 145.38 |
|  | L-cone | 7.62 | 127.07 | 123.24 | 109.81 | 182.42 |
|  | Melano pic | 10.59 | 176.49 | 176.49 | 176.49 | 176.51 |
| α-opic Equivalent Daylight Illuminance (EDI) [lx] | S-cone | 6.11 | 101.95 | 165.32 (62%) | 101.03 (-0.9%) | 146.37 (43%) |
|  | M-cone | 5.78 | 96.45 | 95.29 (-1.2%) | 109.55 (13%) | 111.71 (15%) |
|  | L-cone | 5.23 | 87.25 | 84.62 (-3.01%) | 75.40 (-13%) | 125.26 (43%) |
|  | Melano pic | 8.93 | 148.85 | 148.85 (0.0%) | 148.85 (0.0%) | 148.88 (0.0%) |

### Melatonin suppression of cone-modulated flickering light

We assessed the melatonin area under the curve (AUC) during 2 hours of light exposure after habitual bedtime under different lighting conditions. First, we compared the melatonin AUC under dim control light versus constant background light across all participants to evaluate the biological effect of background light on melatonin suppression. Melatonin suppression was stronger under constant background light at 149 lx_mEDI_ compared to baseline dim light exposure at 9 lx_mEDI_, with moderate evidence supporting this difference (**Bayes factors**, **BF10 = 3.66, Figure 2a, e, i**). Estimates, standard deviations (SDs) and 95% credible intervals (CIs) are shown in **Table 2**.

**Figure 2:**
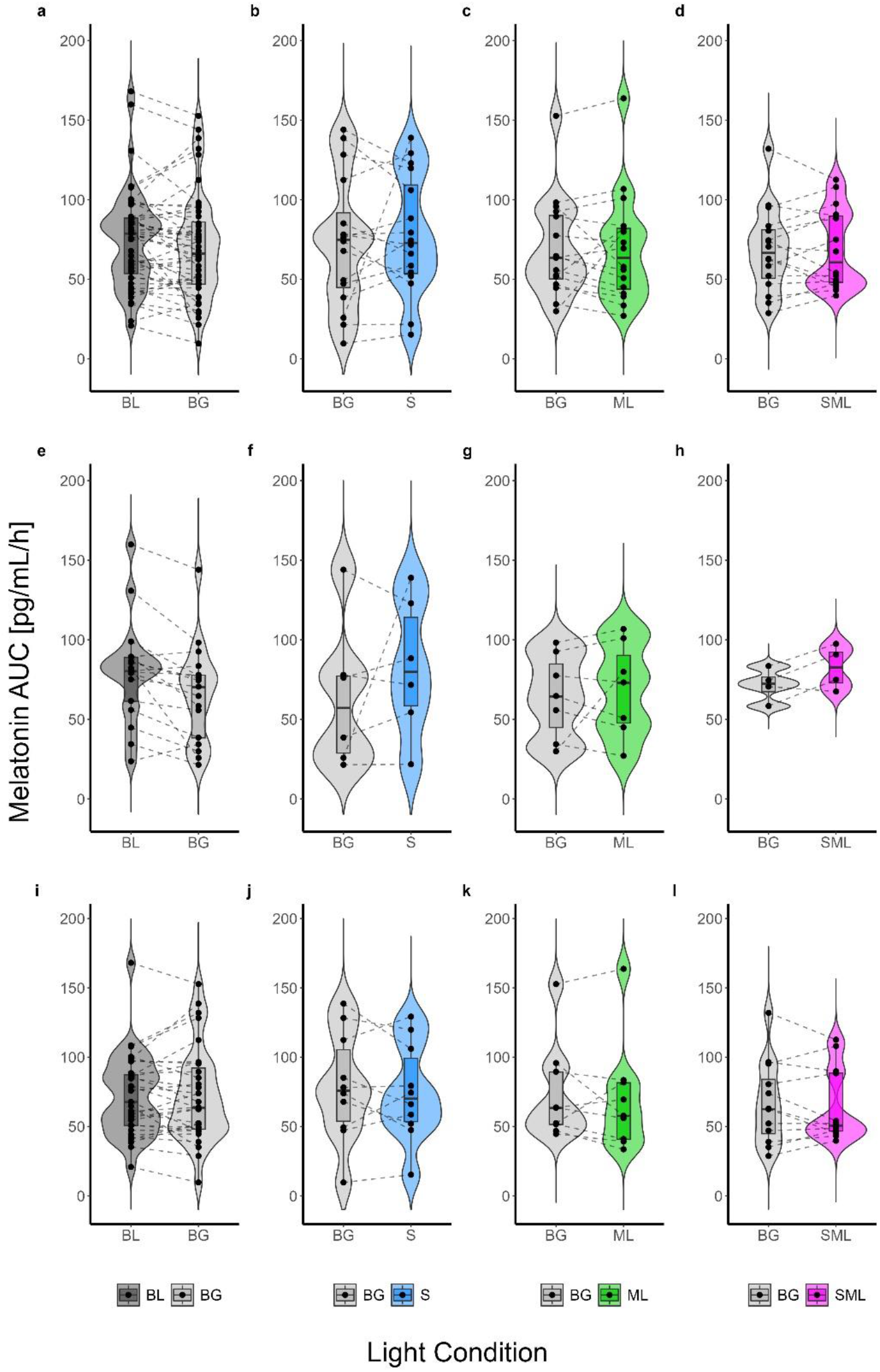
Violin and box plots showing melatonin area under the curve (AUC) [pg/mL/h] during 2 hours of light exposure at night (i.e. the first 2 hours after habitual bedtime) under different light conditions: baseline dim light (BL; dark gray), constant background light (BG; bright gray), and cone-modulated flickering light stimuli targeting S-cones (S; blue), M-L (ML; green), and combined S+M+L-cones (SML; magenta). Panels (a-d) show melatonin AUC comparisons for all participants, while panels (e-h) and (i-l) show melatonin AUC data for winter and summer participants, respectively. (a, e, i) show melatonin AUC for BL and BG conditions. (b, f, j) show melatonin AUC for BG and S-cone flickering light. (c, g, k) compare BG and M-L flickering light. (d, h, l) show melatonin AUC for BG and S+M+L flickering light. Each plot shows individual data points, with the contours of the graphs representing the kernel density estimate of the data distribution. The box plots display the median (central horizontal line), interquartile range (edges of the box, from first quartile; 25^th^ percentile to third quartile; 75^th^ percentile), and the range of minimum and maximum values within 1.5 times the interquartile range from the first and third quartiles (whiskers).

**Table 2:** Melatonin AUC in the dim and background light exposure conditions sampled from the posterior distribution

| Effect | Estimate | SD | 95% CI (lower; upper) |
| --- | --- | --- | --- |
| (intercept) | 2.7 | 2.8 | -2.83, 8.23 |
| Baseline light | 1.7 | 1.8 | -1.92, 5.21 |
| Season | 0.0 | 2.5 | -4.87, 4.88 |
| Baseline light * Season | 3.2 | 2.2 | -1.25, 7.54 |

In contrast, the results for cone-modulated flickering stimuli were inconclusive. For S-cone flickering compared to constant background light, there was anecdotal evidence for melatonin suppression (**BF10 = 0.98, Figure 2b, f, g**). Similarly, M-L cone flickering light stimuli versus background light showed anecdotal evidence for an effect of melatonin suppression (**BF10 = 0.88, Figure 2c, j, k**). Finally, the S+M+L flickering light condition also provided anecdotal evidence compared to constant background light for suppression of melatonin (**BF10 = 0.90, Figure 2d, h, l**). The results of the Bayesian analysis for melatonin AUC for the individual S- cone, M-L and S+M+L groups are shown in **Table 3**.

**Table 3:** Melatonin AUC in the cone-modulated flickering light exposure conditions sampled from the posterior distribution

|  | Effect | Estimate | SD | 95% CI (lower; upper) |
| --- | --- | --- | --- | --- |
| S Group | (intercept) | 1.0 | 3.0 | -4.80, 6.85 |
|  | S-cone light | 0.3 | 2.4 | -4.38, 4.97 |
|  | Season | 0.0 | 2.5 | -4.94, 4.86 |
|  | S-cone light * Season | 0.5 | 2.5 | -4.33, 5.33 |
| M-L Group | (intercept) | 1.5 | 3.0 | -4.49, 7.32 |
|  | M-L light | -0.1 | 2.3 | -4.53, 4.40 |
|  | Season | -0.1 | 2.5 | -4.97, 4.85 |
|  | M-L light * Season | 0.3 | 2.4 | -4.40, 5.05 |
| **S+M+L**<br>**Group** | **(intercept)** | 1.5 | 2.8 | -4.09, 6.89 |
|  | **S+M+L light** | 0.2 | 2.1 | -3.91, 4.43 |
|  | **Season** | 0.1 | 2.5 | -4.79, 4.94 |
|  | **S+M+L light \* Season** | 1.0 | 2.4 | -3.66, 5.75 |

### Subjective alertness of cone-modulated flickering light

We examined subjective alertness using the Karolinska Sleepiness Scale (KSS) during 2 hours of light exposure under different lighting conditions. The results were inconclusive regarding the effect of lighting conditions on alertness. For all participants, the background light did not result in higher alertness compared to the dim light condition (**BF10 = 0.43, Fig. 3a, e, i**). Estimates, standard deviations (SDs) and 95% credible intervals (CIs) for all participants are shown in **Table 4**.

**Figure 3:**
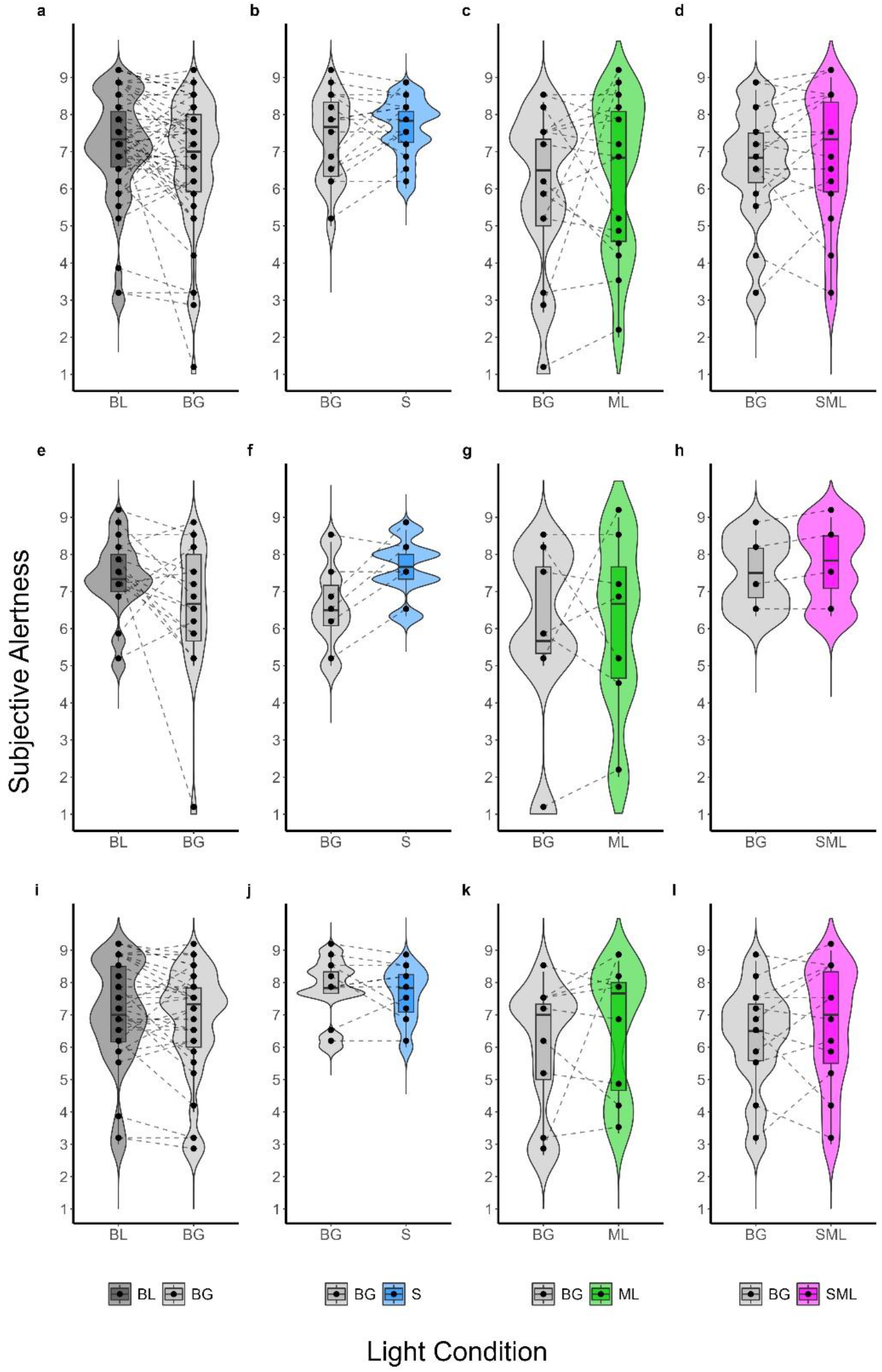
Violin and box plots showing subjective alertness averaged during 2 hours of light exposure at night (i.e. the first 2 hours after habitual bedtime) under different light conditions: baseline dim light (BL; dark gray), constant background light (BG; bright gray), and cone- modulated flickering light stimuli targeting S-cones (S; blue), M-L (ML; green), and combined S+M+L-cones (SML; magenta). Panels (a-d) show average KSS comparisons for all participants, while panels (e-h) and (i-l) show average KSS data for winter and summer participants, respectively. (a, e, i) show average KSS for BL and BG conditions. (b, f, j) show average KSS for BG and S-cone flickering light. (c, g, k) compare BG and M-L flickering light. (d, h, l) show average KSS for BG and S+M+L flickering light. Each plot shows individual data points, with the contours of the graphs representing the kernel density estimate of the data distribution. The box plots display the median (central horizontal line), interquartile range (edges of the box, from first quartile; 25^th^ percentile to third quartile; 75^th^ percentile), and the range of minimum and maximum values within 1.5 times the interquartile range from the first and third quartiles (whiskers).

**Table 4:** KSS subjective alertness in the dim and background light exposure conditions sampled from the posterior distribution

| Effect | Estimate | SD | 95% CI (lower; upper) |
| --- | --- | --- | --- |
| **(intercept)** | 6.6 | 0.3 | 6.06, 7.20 |
| **Baseline light** | 0.3 | 0.3 | -0.23, 0.77 |
| **Season** | -0.2 | 0.5 | -1.12, 0.78 |
| **Baseline light \* Season** | 0.6 | 0.4 | -0.28, 1.39 |

For the S-cone group, the comparison of S-cone flickering light with constant background light showed anecdotal evidence of a difference in subjective alertness (**BF10 = 0.52, Fig. 3b, f, j**). In the M-L group, there was moderate evidence for the null hypothesis, indicating no effect of M-L flickering light on subjective alertness compared to background light (**BF10 = 0.12, Fig. 3c, g, k**). In the S+M+L group, flickering light stimuli provided strong evidence against the null hypothesis, showing no significant difference in alertness compared to constant background light (**BF10 = 0.05, Fig. 3d, h, l**). The results of estimates, SDs, and 95% CIs of subjective alertness in the subgroup analyses for the S-cone, M-L and S+M+L-cone groups are shown in **Table 5**.

**Table 5:** KSS subjective alertness in the cone-modulated flickering light exposure conditions sampled from the posterior distribution

|  | Effect | Estimate | SD | 95% CI (lower; upper) |
| --- | --- | --- | --- | --- |
| **S**<br>**Group** | **(intercept)** | 7.7 | 0.3 | 7.05, 8.27 |
|  | **S-cone light** | -0.1 | 0.3 | -0.71, 0.44 |
|  | **Season** | -1.1 | 0.5 | -2.03, -0.11 |
|  | **S-cone light \* Season** | 1.1 | 0.5 | 0.16, 2.02 |
| **M-L**<br>**Group** | **(intercept)** | 5.8 | 0.7 | 4.33, 7.21 |
|  | **M-L light** | 0.5 | 0.7 | -0.97, 1.98 |
|  | **Season** | -0.2 | 1.0 | -2.20, 1.87 |
|  | **M-L light \* Season** | -0.3 | 1.1 | -2.38, 1.82 |
| **S+M+L**<br>**Group** | **(intercept)** | 6.2 | 0.5 | 5.20, 7.10 |
|  | **S+M+L light** | 0.3 | 0.4 | -0.47, 0.98 |
|  | **Season** | 1.0 | 0.9 | -0.74, 2.84 |
|  | **S+M+L light \* Season** | 0.0 | 0.7 | -1.40, 1.43 |

## Discussion

Despite increasing evidence from non-human models suggesting cone contributions to circadian regulation^26,27,29,49,50^, our results provided no conclusive evidence regarding the role of cone-modulated flickering light in melatonin suppression and subjective alertness in humans with normal trichromatic vision. While previous research^19,20,51,52^ has shown that moderate evening light exposure can suppress melatonin, our study found only moderate suppression under 149 lx_mEDI_ light exposure for 2 hours. In our dataset, seasonal variations have influenced light sensitivity, with higher sensitivity observed in winter^53–55^, possibly due to increased daylight exposure during the summer months^56,57^. Methodologically, we used broadband light exposure without pupil dilation in a setting where participants viewed a normal-size display. This differs from the full-field exposure with pupil dilation used in both the Thapan^19^ and Brainard^20^ studies. Thapan applied 30-minute light pulses shortly after the participant’s habitual bedtime, whereas Brainard’s 1.5-hour exposure began at 2:00 a.m., with no control for individual habitual bedtimes. These critical differences in light exposure duration, spectral composition, and pupil dilation likely contributed to the stronger melatonin suppression effects reported in their studies compared to our findings under the constant background light condition.

Our findings regarding the S-cone contribution are consistent with Spitschan et al^58^ who found no evidence of S-cone contributions to neuroendocrine responses or alertness under evening light exposure prior to individuals’ habitual bedtime. In contrast, Brown et al^59^ reported some evidence for S-cone contribution under short-term (30 min) monochromatic light exposure. The discrepancy may be due to differences in experimental design, including light exposure duration, composition, timing, contrast, and control conditions. In our study, participants were exposed to flickering light stimuli for 2 hours after their habitual bedtime, with 62% S-cone contrast compared to background light, whereas Spitschan’s study used polychromatic light stimuli for 1.5 hours, starting half an hour before participants’ habitual bedtime. Despite a high contrast (8268%) in their study between maximal and minimal S-cone stimulation, similarly to us, they did not find evidence for an S-cone contribution to melatonin suppression. In contrast, Brown’s study used monochromatic light for 30 minutes, focusing on specific wavelengths (415 nm) compared to <10 lx dim light condition. In their study, the S-cone contribution was modelled as a _∼_2:1 ratio with melanopsin signals during short-term light exposure. Six participants were exposed to light timed to the rising phase of the melatonin rhythm, between 23:30 and 01:30. In addition, participants in Brown’s study were given pupil dilators 90 minutes before exposure to light, which may have increased the effects of light.

Recently, Blume et al^60^ also found no conclusive evidence for the contribution of cone photoreceptors to circadian phase delays and melatonin suppression under different flickering light conditions for 1 hour, starting half an hour after habitual bedtime. In their study, participants were exposed to light stimuli designed to activate specific post-receptoral channels along the blue-yellow axis using calibrated silent-substitution methods. These stimuli targeted the S-cone (blue-yellow) and L+M (luminance) pathways to investigate the potential role of colour vision mechanisms in influencing circadian physiology. Despite using both flickering blue-dark and yellow-bright light, all calibrated to ensure equal melanopsin excitation, based on Mouland’s^26^ finding in mice, their results showed no significant differences in circadian phase delays, melatonin suppression, and subjective sleepiness between the different light conditions. The lack of significant findings in their study is consistent with our results, suggesting that even when different post-receptoral channels (i.e. M-L and S+M+L) are selectively activated, cone photoreceptors do not appear to contribute significantly to neuroendocrine responses such as melatonin suppression.

Previous studies by Gooley et al^61^ and St. Hilaire et al^62^, using the same dataset, suggested that cone photoreceptors contribute significantly to the early stages of melatonin suppression and circadian phase resetting during long-term light exposure (6.5 h), particularly at low irradiance levels. Gooley showed that green light (555 nm) was as effective as blue light (460 nm) in suppressing melatonin at the beginning of the exposure, with melanopsin becoming more dominant as the exposure duration increased. Similarly, St. Hilaire extended these findings, showing that S-cones and L+M contributed significantly to melatonin suppression and circadian resetting in the early stages of exposure, with melanopsin taking over in later stages. In contrast, in our study, which used shorter nocturnal light exposures (2 h), inconclusive evidence for a cone contribution was found. These findings suggest that prolonged and continuous light exposure, particularly monochromatic light, may be essential to induce robust cone-driven effects on circadian physiology in humans. However, this setup does not reflect typical lighting conditions in the real world.

In our study, we utilized S+M+L and M-L flickering stimuli designed to target post-receptoral channels known to elicit responses in visual cortical areas, such as V1 and V2/V3, which are critical for visual processing. Previous research^42^ using BOLD fMRI showed higher temporal sensitivity in these areas to S+M+L modulations, although melanopsin-directed flicker did not evoke a comparable response. Despite these visual pathways being sensitive to cone- modulated flickering light, we did not observe significant effects on circadian physiology or melatonin suppression in our study. Another study by Kozaki et al^41^, demonstrated that flickering light can enhance melatonin suppression. Their study found that 100 Hz flickering blue light was more effective at suppressing melatonin than non-flickering blue light. However, unlike Kozaki et al, our use of flickering stimuli targeting cone photoreceptors did not result in a greater degree of melatonin suppression, potentially due to differences in flickering frequencies. As shown by Brainard et al^63^, intermittent light exposure (10-minute pulses) leads to less melatonin suppression compared to continuous exposure. In addition, Kozaki’s study used blue light (465 nm), which is likely to activate melanopsin in addition to cones, as they did not specifically isolate cone-mediated responses. This difference may explain why we did not observe the same degree of melatonin suppression in our study.

The absence of a conclusive evidence for cone involvement in melatonin suppression and light’s alerting response raises the possibility that post-receptoral channels may not play a physiologically significant role in the circadian system under the specific conditions examined. It appears that light-responsive neuroendocrine pathways are predominately tuned to signals from melanopsin-expressing cells rather than to cone inputs, which may function primarily within the visual system.

## Conclusion

Our results suggest that S-cones, M-L and S+M+L cone combinations do not contribute significantly to the melatonin suppression and alerting response to nocturnal light exposure that we typically experience during screen work. These findings may help in the design of display screen lighting systems to minimise the impact on the circadian physiology of night workers.

## Study Limitations

While our study aimed to address several key aspects, some limitations should be acknowledged. First, pupil size was not measured steady-state across conditions for all participants. Variations in pupil size may have influenced retinal irradiance under cone- modulated flickering light conditions. Although this could have affected light exposure at the retina, it is unlikely that small differences in retinal irradiance would have significantly masked true photoreceptor-mediated effects. Recent evidence^64^ suggests that higher mEDI is associated with smaller pupil sizes. However, in our study, the mEDI was constant between the cone flickering and background light conditions. Furthermore, while we assumed that rods did not contribute to circadian responses under photopic conditions, recent evidence^65^ suggests that rod inputs may play a role in circadian photoentrainment at higher light levels. Finally, this study focused on the short-term effects of light exposure, and the potential long-term effects of cone-modulated light remain unexplored. Future research should include studies conducted in more ecologically valid settings, with larger and more diverse participant groups, and consider varying light exposure conditions to further elucidate the role of cone photoreceptors in circadian and neuroendocrine regulation.

## Method

### Participants

From 85 interested participants, 48 healthy young adults (50% female) aged 18–35 years (mean age = 25.11 ± 4.4 years) with normal colour vision completed the study protocol over one year of data collection, starting from October 2022 and ending in October 2023, at the Centre for Chronobiology in Basel, Switzerland. After an initial telephone interview, screening was carried out in three phases: an online phase (REDCap^66^), an in-person phase, and final assessments by a physician and an ophthalmologist. Exclusion criteria included body mass index (BMI) <18.5 or >29.9, use of chronic medications affecting the neuroendocrine, sleep, circadian physiology, or visual systems, and use of drugs or nicotine. Participants were also excluded if they had worked shift work in the previous 3 months or travelled across more than two time zones in the month prior to the study. Other exclusion criteria were high myopia or hyperopia (>-6 or <6 diopters), photosensitive epilepsy, and any ocular disease. Female participants were also excluded if they were pregnant, breastfeeding or using hormonal contraceptives. Further details on participant demographics, exclusion criteria, and number of withdrawals are provided in **Supplementary Table 2, Table 3, and Figure 1**. All participants received financial compensation for their participation.

### Colour vision assessment

Colour vision was assessed at several stages of the screening process. During the online screening, participants completed the Ishihara test^67^. At the in-person screening, participants underwent the Cambridge Colour Test (CCT)^68,69^ and the Farnsworth-Munsell 100 Hue Test^70,71^, with assessment times recorded for each. Finally, during the ophthalmological screening at the University Hospital Basel, a comprehensive colour vision assessment was performed by an ophthalmologist (F.Y.). Exclusion criteria are provided in **Supplementary Table 3**.

### Light setting

We used a modified monitor equipped with five LED primaries^72^ (430 nm, 480 nm, 500 nm, 540 nm, and 630 nm), with calibration performed at eye level using a JETI spectroradiometer 1501 (JETI Technische Instrumente GmbH, Jena, Germany). Calibration was performed with the monitor positioned at a height of 120 cm, at a distance of 65 cm from the participant, and tilted downward by 20 degrees. Participants were exposed to three different lighting conditions: (1) a constant low-lux control condition (BL; 8 lx, 9 lx melanopic equivalent daylight illuminance [mEDI]), (2) a constant background light (BG; 100 lx, 149 lx mEDI), and (3) cone-modulated flickering light targeting different cone combinations and post-receptoral channels. The cone-modulated stimuli were as follows: S-cones, with no change in L, M or melanopsin activation (42 lx, 149 lx mEDI, 62% S-contrast), M-L, with no change in S-cone or melanopsin activation (96 lx, 149 lx mEDI, 13% M-contrast, -13% L-contrast) and S+M+L, with no change in melanopsin activation (167 lx, 149 lx mEDI, 43% S-, 15% M- and 43% L-contrast). We measured the spectral shifts of the primaries in 16 settings, each with 8-bit resolution. In the control dim light condition, approximately 3% of each primary was activated, while in the background light condition, 50% of each primary was used. During the background condition, the light was designed to maintain a constant photoreceptor activation profile, while during the modulation condition, the light flickered sinusoidally at 1 Hz (30 seconds on, 30 seconds off).

### Study design

This study was a randomised within-subject design with a protocol consisting of 9-hour laboratory visits in constant environmental conditions. The study procedures were identical for all participants and scheduled according to the participants’ habitual bedtime (HBT) starting 6 hours before to 3 hours after their HBT. The only difference was a 2 h light intervention comprised of dim, constant background and cone-modulated flickering light. All participants were exposed to dim and constant light and one-third were exposed to either S- cone, M-L, or S+M+L light conditions. During the control evening, participants were exposed to dim light for 7.5 hours, which served as the dim light visit. In the background evening, participants were exposed to dim light for the first 5 hours, followed by 2 hours of constant illumination after HBT. During the cone-modulated evening, the same as the background, they were first under dim light and then under cone-modulated flickering light for 2 hours after their HBT. Participants were instructed to keep their eyes open and fixate on the centre of the display during the 2 hours of light exposure, using a chin rest. They also underwent a 20- minute period of complete dark adaptation before exposure to the light conditions. In addition, two pupillometry sessions of 45 minutes each were conducted: one prior to dark adaptation and one after light exposure (i.e. the results will be published elsewhere). One week before the study session, participants underwent a three-week ambulatory phase, refraining from alcohol, irregular caffeine intake, and on the day of the study session refraining from bananas, and citrus fruits to avoid potential impacts on salivary melatonin^73^. They were instructed to maintain a regular sleep-wake rhythm, monitored through wrist actimetry (i.e. Condor Instruments ActTrust 2) and an online sleep diary questionnaire in REDCap^74,75^. Participants deviating by more than 30 minutes in bedtime or wake time once within the five days preceding each experimental visit were postponed or excluded. One day before the experimental sessions, participants refrained from intense exercise to maintain constant physical exhaustion^76^. Upon arrival to the study sessions, participants were screened for alcohol (i.e. ACE X Alcohol test; ACE Instruments, *Germany*) and urine multi-drug (i.e. Drug-Screen-Multi 6; nal von Minden, Den Haag, The Netherlands) testing. Saliva samples were collected every 30 minutes, and subjective alertness ratings^77^ (KSS) were gathered hourly throughout the sessions. During the laboratory sessions, participants were not allowed to use mobile phones or laptops and were kept unaware of external time cues. Reading, solving puzzles, or listening to podcasts were allowed, provided they did not involve self-luminous displays. Participants were asked to maintain a consistent level of activity across sessions. To control for potential masking effects, participants received identical meals consisting of three sandwiches at 1.5, 5.5 and 8.5 hours after arrival. They had constant access to water, and dark sunglasses were provided during toilet breaks to minimise unwanted light exposure. As the study ended after midnight, participants were allowed to sleep at the Centre for Chronobiology after the sessions.

### Salivary melatonin

Saliva samples were collected at 30-minute intervals throughout the study session, from 6 hours before the participants’ habitual bedtime to 3 hours after. Melatonin concentrations were quantified using a direct double antibody radioimmunoassay (RK-DSM2)^78^. The analytical sensitivity of this assay was 0.2 pg/mL, with a limit of quantification of 0.9 pg/mL in saliva. Measurements were performed by NovoLytiX GmbH, Witterswil, Switzerland, according to the manufacturer’s instructions for the RK-DSM2 test kit (version 5, 2023-06-08). The area under the curve (AUC) for melatonin during light exposure was calculated using the trapezoidal method.

### Subjective alertness

Subjective alertness was assessed using the 9-point Karolinska Sleepiness Scale (KSS)^79^, which was administered hourly during the study (a total of 10 assessments per session). To assess sleepiness during light exposure, the KSS scores for each participant were averaged over the 2 hours of light exposure period under each light condition.

### Statistics and reproducibility

A Bayesian approach was used for all statistical analyses using R (version 4.3.0, R Core Team, 2023) with the packages rstanarm^80,81^ and bayestestR^82^. Priors were defined as normally distributed with a mean of 0 and a standard deviation (SD) of 1. Model estimation was performed using four chains of 20,000 iterations each to ensure accurate estimation of Bayes factors and credible intervals. We report 95% credible intervals for all model estimates. The analysis focused on modelling the area under the curve (AUC) for melatonin levels and average subjective alertness (KSS scores) during the 2-hour light exposure period. Based on our ongoing research indicating a significant seasonal effect on melatonin suppression, we included the interaction of light stimuli and season in our analysis. Both null (without light condition) and alternative models (with light condition) were compared using Bayes factors to assess the effect of light conditions.

## Acknowledgements

We are grateful to all the participants in this project. The valuable contributions of our interns and study helpers, including Sonja Camenzind, Rebecca Frommherz, Maria Vettiger, Fiona Vogel, Lisa Tran, Taifun Süner, Nadja Tschan, Bela Bernasconi, Milica Trailović, Karoline Edrich, Melanie Schmid and Joshua Reese, were instrumental in the success of this research. We also thank Jakob Weber and Alejandro Fernandez Estrada from NovoLytiX for their assistance with the melatonin assay, and Christine Blume for providing access to equipment and laboratory facilities.

## Author contributions

Conceptualization, F.F., R.L., M.S., O.S., and C.C.; Methodology, F.F., M.S., O.S., and C.C.; Validation, F.F.; Formal Analysis, F.F.; Data Curation, F.F.; Investigation, F.F., F.Y., and C.E.; Visualization, F.F. and C.C.; Funding Acquisition, C.C.; Resources, C.C.; Supervision, F.F., M.S., O.S., and C.C.; Project Administration F.F. and C.C.; Writing – Original Draft, F.F.; Writing – Review & Editing, R.L., F.Y., C.E., M.S., O.S., and C.C.

## Declaration of generative AI and AI-assisted technologies in the writing process

During the preparation of this work, the author(s) used ChatGPT -4o and DeepL Write to enhance the clarity, coherence, and overall quality of the text. After using this tool or service, the author(s) reviewed and edited the content as needed and take(s) full responsibility for the content of the publication.

## Supplemental information titles and legends

Table S1- S3 and Figure S1

## Ethics Approval Statement

The study received ethical approval from the Ethics Commission of Northwest and Central Switzerland (EKNZ) under approval number 2022-00401, classified as an "other clinical trial". All procedures were performed in accordance with the principles of the Declaration of Helsinki. Participants received detailed information about the study and gave written informed consent before participating. All participants were offered compensation for their time and participation in the study.

## Clinical Trial Registration

This study was conducted according to the protocol ID 2022-00401. The trial was registered at ClinicalTrials.gov with the identifier NCT05423002.

## Resource availability Lead contact

Further information and requests for resources and reagents should be directed to and will be fulfilled by the lead contact, Christian Cajochen.

## Materials availability

The code used for generating the light stimuli in this study will be made available upon request. However, we may require a payment and/or a completed materials transfer agreement if there is potential for commercial application, as we plan to explore commercial opportunities in the future.

## Data and code availability

De-identified human data and laboratory log have been deposited on the FigShare as [Data: https://figshare.com/s/fb65e5617d0c783c9e94] and [Laboratory log: https://figshare.com/s/35058f9443d2478f05ee] are publicly available as of the date of publication.

All original code has been deposited in the GitHub repository and is publicly available at [https://github.com/mahsafazlali/cone-photoreceptors-contribution] as of the date of publication.

Any additional information required to reanalyze the data reported in this paper is available from the lead contact upon request.

