## Supplementary material for "Lack of evidence for the contribution of cone photoreceptors to human melatonin suppression and alerting response to light at night": Table S1, Table S2, Table S3, Figure S1: FazlaliEtAl_Supplementary Material.docx

Table S1) The Bayes Factor (BF) interpretation based on Jeffreys (1961)

| Bayes Factor | Interpretation |
| --- | --- |
| > 100 | Decisive evidence for H1 |
| 30 - 100 | Very strong evidence for H1 |
| 10 - 30 | Strong evidence for H1 |
| 3 - 10 | Moderate evidence for H1 |
| 1 - 3 | Anecdotal evidence for H1 |
| 1 | No evidence |
| 1/3 - 1 | Anecdotal evidence for H0 |
| 1/10 – 1/3 | Moderate evidence for H0 |
| 1/30 - 1/10 | Strong evidence for H0 |
| 1/100 - 1/30 | Very strong evidence for H0 |
| < 1/100 | Decisive evidence for H0 |

Table S2) participants' characteristics in different cone-modulated flickering groups. Data are expressed as mean (± SD). BMI: Body Mass Index; BDI-II: Beck Depression Inventory; µMCTQ: Ultra-Short Version of the Munich ChronoType Questionnaire; PSQI: Pittsburgh Sleep Questionnaire; ESS: Epworth Sleepiness Scale; Visual Acuity with single Landolt C; Farnsworth Munsell 100 Hue test; Cambridge Colour Test Trivector; Protan, Deutan, and Tritan

| **Descriptives** | | **Group 1 (S-Cone)** | **Group 2 (M-L)** | **Group 3 (S+M+L)** |
| --- | --- | --- | --- | --- |
| **Age** | | 25.72 (3.74) | 26.11 (5.08) | 23.50 (4.17) |
| **BMI** | | 22.10 (2.44) | 22.31 (2.87) | 22.93 (1.61) |
| **BDI_II** | | 2.38 (2.60) | 2.13 (2.55) | 4.63 (4.75) |
| **µMCTQ** | | 4.38 (0.70) | 3.98 (0.83) | 3.95 (0.91) |
| **PSQI** | | 3.31 (1.20) | 2.88 (1.75) | 3.44 (1.36) |
| **ESS** | | 6.31 (3.32) | 5.25 (2.86) | 5.63 (2.73) |
| **Visual Acuity** | | 1.96 (0.44) | 1.78 (0.42) | 1.99 (0.32) |
| **Farnsworth** | | 12.38 (3.88) | 13.38 (4.43) | 13.38 (5.45) |
| **CCT** | **Protan** | 4.73 (1.79) | 5.13 (1.56) | 4.08 (1.37) |
|  | **Deutan** | 4.77 (1.61) | 4.66 (1.55) | 4.57 (1.69) |
|  | **Tritan** | 6.70 (2.79) | 6.23 (2.30) | 6.24 (2.64) |

Table S3) inclusion and exclusion criteria of online, in-person, physician and ophthalmology, and every session screening

|  | **Aspect** | **Assessment modality** | **Exclusion criterion and cut-off** |
| --- | --- | --- | --- |
| **Online screening** | Age | Self-report | <18 years and >35 years |
|  | BMI | Self-reported height and weight | <18.5 and >29.9 |
|  | Pregnancy (only female) | Self-report | 'Yes' response |
|  | Use of hormonal contraceptives (only female) | Self-report | 'Yes' response |
|  | Lactation or breastfeeding (only female) | Self-report | 'Yes' response |
|  | Menstrual cycle (only female) | Reproductive Status Questionnaire for Menstrual Cycle Studies (ref) |  |
|  | Color vision deficiency (only for category 2) | Ishihara Test (r | ' Yes ' response |
|  | Chronotype | Ultra-short Munich Chronotype Questionnaire (µMCTQ) | ≤ 2 and ≥7 |
|  | Sleep duration | Ultra-short Munich Chronotype Questionnaire (µMCTQ) | < 6 and > 9 |
|  | Sleep quality | Pittsburgh Sleep Quality Index, PSQI | >5 |
|  | Smoking | Self-report | >0 |
|  | Substance abuse | Alcohol Use Disorders Identification Test, AUDIT | >7 |
|  | Depressive symptoms | BDI-II | >13 |
|  | High myopia | Self-report from prescription information | < -6 diopters |
|  | High hyperopia | Self-report from prescription information | > +6 diopters |
|  | Transmeridian travel (>2 zones) <1 month prior to first session | Self-report | 'Yes' response |
|  | Shift work <3 months prior to study | Self-report | 'Yes' response |
|  | Current participation in other clinical trials | Self-report | 'Yes' response |
|  | Any ophthalmological or optometric conditions (cataract, glaucoma, retinal detachment, macular conditions, chronic inflammations, eye injuries or operations) | Self-report | Any 'Yes' response |
|  | Any general health concerns or disorders, including heart and cardiovascular, neurological, nephrological, endocrinological and psychiatric conditions | Self-report | Any 'Yes' response |
|  | Any chronic medication affect on sleep | Self-report | Any 'Yes' response |
| **in-person screening** | BMI | Measured height and weight | <18.5 and >29.9 |
|  | Pregnancy test (only women) | M-Budget pregnancy test | Positive test |
|  | Normal color vision | Cambridge Color Test, Farnsworth Munsell 100 Hue Test | Protan >10, Detran > 10, Tritan > 15, and 100 Hue score > 40 |
|  | Normal best-corrected visual acuity (BCVA) | Landolt C test | Visus < 0.5 |
|  | Ability to understand study language | In-person interaction with the experimenter | Experimenter judgment |
| **Physician and ophthalomology screening** | Color vision deficiency | Check by ophthalmologist | Ophthalmologist judgment |
|  | Risk of angle-closure glaucoma | Check by ophthalmologist | Ophthalmologist judgment |
|  | Any ophthalmological or optometric conditions (cataract, glaucoma, retinal detachment, macular conditions, chronic inflammations, eye injuries or operations) | Check by ophthalmologist | Ophthalmologist judgment |
|  | Any general health concerns or disorders, including heart and cardiovascular, neurological, nephrological, endocrinological and psychiatric conditions | Check by physician | Physician judgment |
|  | Any chronic medication affect on sleep | Check by physician | Physician judgment |
| **Every session screening** | Drug use (AMP, BZD, COC, MOR/OPI, MTD and THC) | Drug-Screen-Multi 6; nal von Minden | Any positive test |
|  | Alcohol use | Breathalyzer ACE X | >0.05 |
|  | Sleep-wake times in 5 days prior to each experimental session | Actigraphy record and sleep diary | >1 deviation from ±30 minute window sleep and wake-up time |
|  | Ability to follow study instructions | In-person interaction with the experimenter | Experimenter judgment |

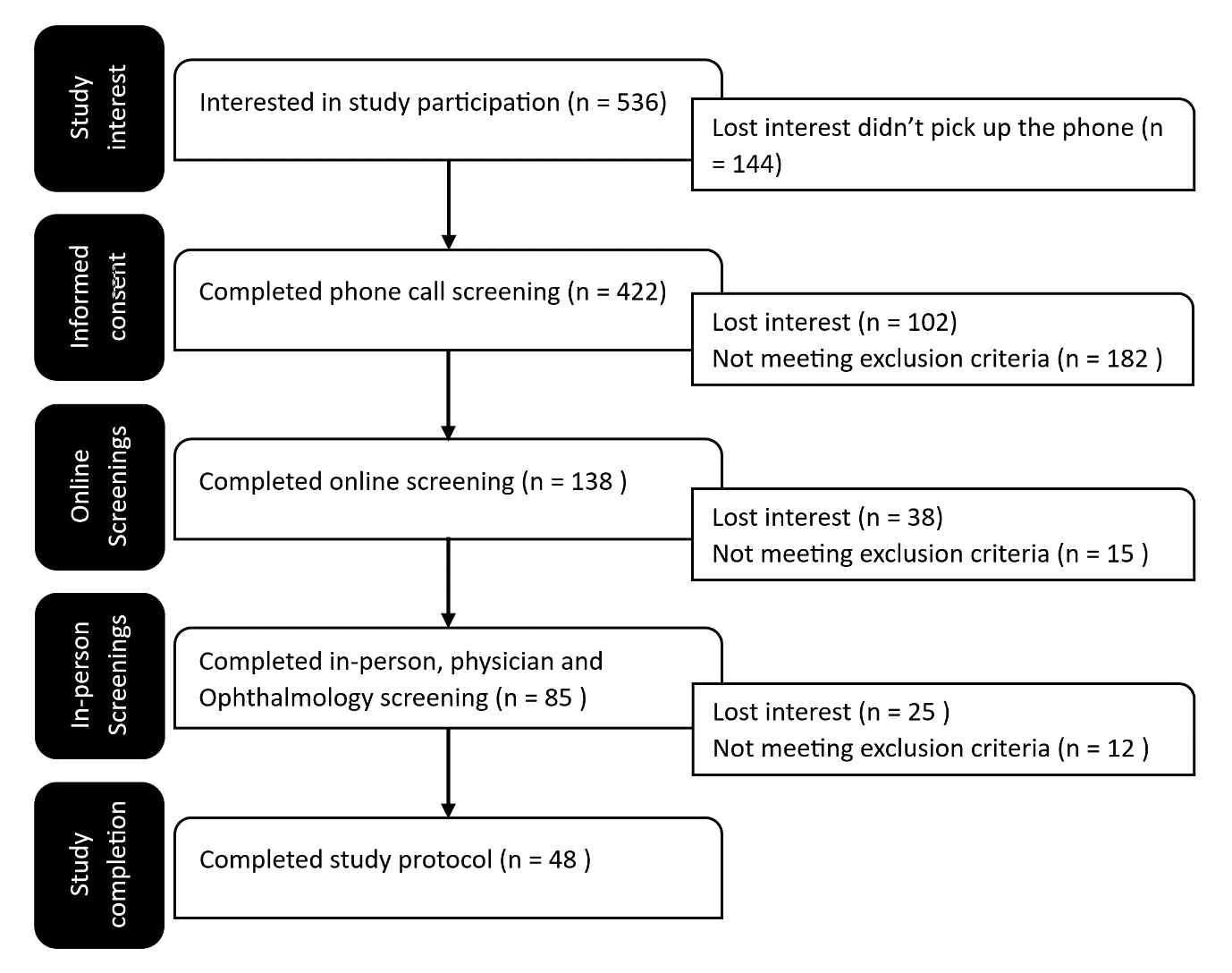

Figure S1) Study flowchart from initial interest to completion the study sessions. A total of 536 volunteers initially expressed interest in participating in the study, of whom 422 attended the mandatory telephone information session. Of these, 138 participants signed the informed consent form and completed the online screening. Of these, 85 attended the in-person, physician and ophthalmological screening. During the process, 25 participants lost interest, 7 were excluded due to non-compliance with the agreed sleep-wake schedule, 2 were excluded due to a positive drug test, and 3 withdrew after the first study session. In the end, 48 participants completed the study.
